# Evaluating Quantitative and Functional MRI As Potential Techniques to Identify the Subdivisions in the Human Lateral Geniculate Nucleus

**DOI:** 10.1101/2022.11.16.516765

**Authors:** Irem Yildirim, Khan Hekmatyar, Keith A. Schneider

**Author notes:** Author contributions. IY: Conceptualization, Formal analysis, Investigation, Methodology, Software, Visualization, Writing - original draft, Writing - review & editing; KH: Methodology, Resources; KAS: Conceptualization, Methodology, Software, Writing - review & editing, Supervision, Funding acquisition.

## Abstract

Segmenting the magnocellular (M) and parvocellular (P) divisions of the human lateral geniculate nucleus (LGN) has been challenging yet remains an important goal because the LGN is the only place in the brain where these two information streams are spatially disjoint and can be studied independently. Previous research used the amplitude of responses to different types of stimuli to separate M and P regions (Denison et al., 2014; Zhang et al., 2015). However, this method is confounded because the hilum region of the LGN exhibits greater response amplitudes to all stimuli and can be mistaken for the M subdivision (DeSimone & Schneider, 2019). Therefore, we have employed two independent methodologies that do not rely upon the functional response properties of the M and P neurons to segment the M and P regions: 1) structural quantitative MRI (qMRI) at 3T to measure the T1 relaxation time, and 2) monocular and dichoptic functional MRI (fMRI) procedures to measure eye-specific responses. Our qMRI results agreed with the anatomical expectations, identifying M regions on the ventromedial surface of the LGN. The monocular fMRI procedure was better than the dichoptic condition to identify the eye-dominance signals. Both procedures revealed significant right eye bias, and neither could reliably identify the first M layer of the LGN. These findings indicated that the qMRI methods are promising whereas the functional identification of contralateral layers requires further refinement.

**Highlights:**

- T1 parameter in qMRI segregates M and P regions of LGN in individual subjects at 3T.
- Eye-specific voxels in LGN respond more strongly to monocular than dichoptic viewing.
- Clusters of eye-specific regions but not layers can be separated at 1.5 mm resolution.

## 1. Introduction

The LGN is the visual relay in the thalamus (Nassi & Callaway, 2009; Skalicky, 2016). It receives projections from retinal ganglion cells, projects primarily to the primary visual cortex (V1), and also receives massive feedback from V1. It has a laminar structure, with typically six monocular layers in humans, receiving input alternatingly from the contralateral or ipsilateral eye (Figure 1a). Specifically, Layers 2, 3, and 5 consist of ipsilateral eye neurons while Layers 1, 4, and 6 are contralateral eye neurons. The four dorsal layers (Layers 3 to 6) are composed of parvocellular (P) neurons while the two ventral layers (Layers 1 and 2) are composed of magnocellular (M) neurons (Figure 1b), receiving input from the midget and parasol ganglion cells in the retina respectively. The M and P neurons in LGN differ in their functional roles (Maunsell, 1992; Merigan & Maunsell, 1993). The M neurons are specialized to encode coarse and transient characteristics, such as luminance (Shapley & Perry, 1986) and temporal frequency of a signal (Derrington & Lennie, 1984), while the P neurons encode detailed and sustained characteristics such as color and form (Livingstone & Hubel, 1988).

**Figure 1.**
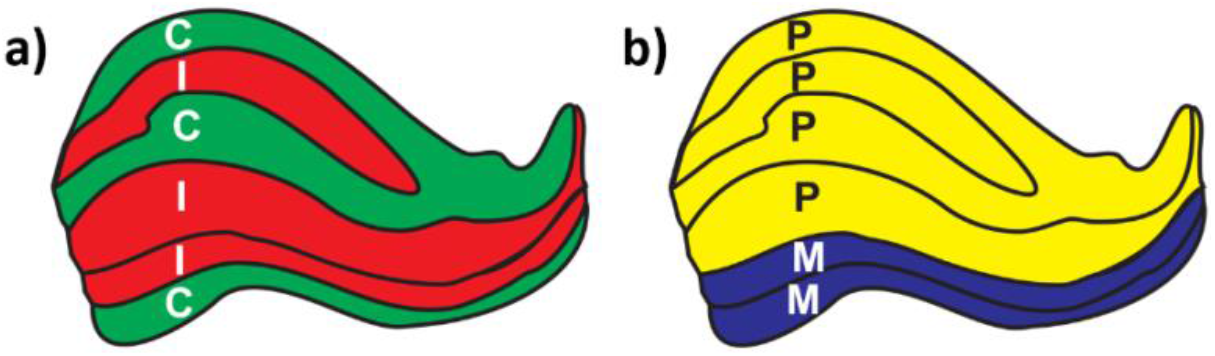
The structure of the lateral geniculate nucleus. M = Magnocellular, P = Parvocellular, C = Contralateral, I = Ipsilateral. Tracings were generated based on Andrews et al. (1997).

The M and P pathways are of considerable interest for their roles in the mechanisms of visual perception and consciousness (Breitmeyer, 2014; Denison & Silver, 2012; Milner, 2012) and in clinical disorders such as dyslexia (Stein, 2001; Stein & Walsh, 1997) and schizophrenia (e.g., Butler & Javitt, 2005; Schechter et al., 2003). Studying these pathways independently, however, has been challenging due to the intermixing of the two pathways starting in V1 (Aleci & Belcastro, 2016; Merigan & Maunsell, 1993). In the LGN, the M and P neurons are completely segregated in separate layers, but the small size of the LGN, with layers on the order of 1 mm thick, approaches the resolution limits of human neuroimaging. Previous MRI attempts have identified the regions at the group level and/or using a group-level criteria such as for the proportion of the M and P sections in LGN (Denison et al., 2014; P. Zhang et al., 2015), but this does not enable the measurement of the properties of the M and P layers in individuals. Further, previous studies, such as Denison et al. (2014), Qian et al. (2020), and Zhang et al. (2015) attempted to identify the M and P regions with fMRI using visual stimuli tuned to the M or P neurons. However, DeSimone and Schneider (2019) showed that the hilum region of the LGN, a vascular region rich with blood vessels and nerves, had larger responses across the range of stimuli. They found that any method based only on the response amplitudes without proper normalization would be likely to mistake the hilum for the M subdivision.

Our aim was to identify the M and P layers of the LGN in individual subjects using anatomical and functional procedures that did not rely upon their differences in functional response properties. Using fMRI, we attempted to segment the contralateral layers of LGN from the bordering ipsilateral layers (Figure 1a), i.e., isolating the ipsilateral cluster of Layers 2 and 3 that separates the contralateral eye layer 1 (M) from Layers 4–6 (P). Previously, Haynes et al. (2005) used a 3T MRI scanner and monocular visual stimulation—participants closed one eye— to distinguish left from right eye signals. More recently, Qian et al. (2020) used a 7T scanner and presented monocular stimuli dichoptically using a fast refresh-rate projector alternating each eye between a visual stimulus while the image to the other eye was blank. They identified two clusters in the LGN, one lateral contralateral and another medial ipsilateral, but not individual layers. Our goal was to compare the monocular and dichoptic viewing conditions in segregating the eye-specific regions in the LGN at 3T in individual subjects, and to determine the possibility of identifying the contralateral M layer.

We also sought to compare the fMRI results to structural methods. Recent developments in quantitative MRI (qMRI) permit measurement of the microstructure of tissues such as myelination (Lutti et al., 2014; Mezer et al., 2013), which can differentiate the M and P regions. The morphology of the M and P neurons in LGN differ, with P neurons having smaller somas and thinner axons and M neurons having larger somas and thicker axons. The density of the P neurons is therefore higher (Andrews & Purves, 1997; Hassler, 1966; Nassi & Callaway, 2009). It is unclear whether this higher density would result in greater overall myelination in the P region (Pistorio et al., 2006), or whether the thicker, more highly myelinated M axons would result in higher overall myelination in the M region (Yoonessi & Yoonessi, 2011). Müller-Axt et al. (2021) recently demonstrated qMRI results in the human LGN consistent with the former. At 7T, they measured shorter T1 relaxation times in the P compared to M regions. Our overall aim was to replicate and compare these different techniques at 3T in individual subjects and to quantify the optimal duration of data acquisition necessary.

## 2. Methods

### 2.1. Participants

This study was approved by the University of Delaware Institutional Review Board. Three healthy participants (aged 28–33 years, 1 male) with normal or corrected-to-normal vision provided written informed consent and were compensated $20/hour.

### 2.2. MRI Procedures and Processing

Each participant was scanned on seven different days (four structural scanning sessions and three functional) for approximately 90 min each day. MRI data were acquired on a 3T Siemens Magnetom Prisma MRI scanner with a 64-channel head coil. We used FSL software (https://fsl.fmrib.ox.ac.uk/fsl/fslwiki/FSL) to process all the MRI data unless otherwise noted. All the raw data is publicly available at https://openneuro.org/datasets/ds004187, the processed data and code are available at https://github.com/yirem/LGN_layers.

#### 2.2.1. T1-weighted MRI

At the beginning of each scanning session, we acquired a 3D MPRAGE sequence (0.7 mm isotropic voxels, repetition time (TR) = 2080 ms, echo time (TE) = 4.64 ms, inversion time (TI) = 1050 ms, flip angle (α) = 9°, field of view (FoV) = 210 mm, phase-encoding acceleration factor = 2, scan time approximately 6 min). All subsequent scans were aligned to this T1-weighted image of each subject and analyzed in their native space.

#### 2.2.2. Quantitative MRI

qMRI data were acquired with a 3D MP2RAGE sequence (0.7 mm isotropic voxels, TR = 5000 ms, TE = 3.6 ms, partial phase Fourier in slice = 6/8, approximately 16 min acquisition time). The sequence had two inversion times and flip angles (TI_1_ = 900 ms, TI_2_ = 2750 ms, α_1_ = 3°, α_2_ = 5°), enabling the calculation of the T1 relaxation time, i.e., the qT1 map. Seventeen scans were acquired for each participant during four sessions on different days, during which participants watched a movie of their choice.

The MP2RAGE sequence simultaneously acquired the T1-weighted (GRE_TI1_) and proton density weighted (GRE_TI2_) image volumes. The uniform T1-weighted image volume was obtained from the real component of the normalized complex ratio from the two acquired image volumes. This process amplifies the noise in the uniform T1-weighted image. Using the MP2RAGE toolbox (https://github.com/benoitberanger/mp2rage) in SPM (https://www.fil.ion.ucl.ac.uk/spm/software/spm12/) in MATLAB (The Math Works, Inc.), the numeric instability in background noise was suppressed by introducing a constant real number (beta) to the uniform T1-weighted image volume (O’Brien et al., 2014). We then computed qT1 maps (also available at https://openneuro.org/datasets/ds004187), a measurement of the T1 relaxation constant for each voxel. qT1 maps were computed for each of the 17 scans for each participant, 16 of which were aligned to the first one.

##### 2.2.2.1. Anatomical LGN Masks

To create the LGN masks, we averaged the 16 qT1 maps. The average qT1 map for each subject was then resampled to double the resolution (0.35 mm isotropic voxels) with a sinc interpolation to reduce partial volume effects. Using this upsampled average qT1 map, we manually masked each LGN for each participant. A typical slice with the LGN outlined is shown for one subject in Figure 2. Care was taken to avoid incorporating the bright (high qT1) cerebrospinal fluid (CSF) into the LGN region of interest (ROI), which would confound the comparison of the M and P regions. This was aided by a darker region located between the CSF and LGN (see Figure 2). The qT1 map and LGN mask were then aligned to each subject’s T1 volume, using 6 degrees of freedom and mutual info as the cost in FSL (Figure 4a).

**Figure 2.**
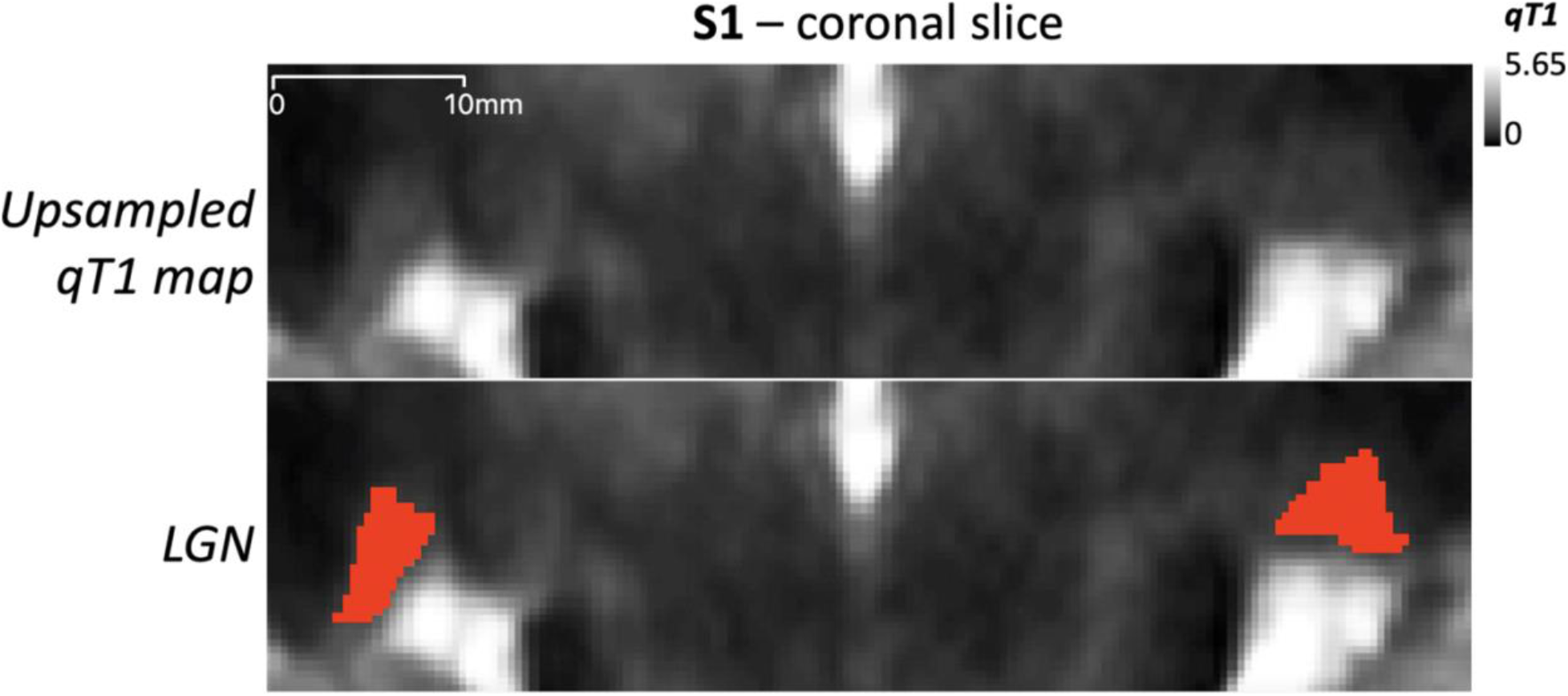
LGN mask on the coronal slice for a representative participant (S1). The upper panel is the upsampled average qT1 map (0.35 mm isotropic voxel resolution), with brighter colors indicating higher qT1 values. In the lower panel, the LGN are outlined in red.

##### 2.2.2.2. qT1 Analysis

The left and right LGN were masked on the average qT1 map for each participant (Figure 4b) and analyzed separately. We expected two distributions of qT1 values within the LGN voxels, corresponding to the M and P sections (Müller-Axt et al., 2021) and therefore fit each qT1distribution to a mixture of two Gaussians using the fitgm function in MATLAB, as Müller-Axt et al. (2021) did with their 7T data. Figure 5a shows the histograms of qT1 maps with the fitted Gaussians. To ensure the two-component model was warranted, we compared the fit of the 2-component Gaussian model to a 1-component Gaussian model for each LGN using both Akaike Information Criteria (AIC) and Bayesian Information Criteria (BIC) using log-likelihoods with a maximum number of 1000 iterations. Every case favored the 2-component model (see Table S1 in Supplemental Materials). Diverging from the methods of Müller-Axt et al. (2021), we then calculated a qT1 threshold between the M and P distributions using the fraction of M voxels to the right of the threshold (dark blue line in Figure 5a):

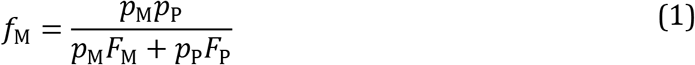

where *p*_M_ and *p*_P_ are the M and P proportions estimated by the Gaussian models, respectively (always arranged such that *p*_M_ < *p*_P_), and *F*_M_ and *F*_P_ are the cumulative distribution functions of the fitted Gaussian distribution for each component:

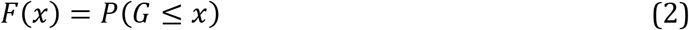

where *x* is the qT1 separation threshold (*x*-axis in Figure 5a) and *P* is the probability that the fitted Gaussian distribution *G* is less than or equal to *x*. We determined the threshold qT1 value (dashed line in Figure 5a) where at least 50% of the voxels to the right belonged to the M distribution. In post-mortem LGN (Müller-Axt et al., 2021), a more pronounced M distribution was evident, and thus, we decided on this liberal threshold because the M voxels were underestimated in the mixture Gaussian model with our 3T data.

##### 2.2.2.3. Random Subsampling Analysis

To determine the number of qT1 maps that needed to be averaged to obtain reliable results, we reran the analysis for each of the 16 individual qT1 maps and for the 16 random subsamples of different sizes, i.e., two to 15 qT1 maps in a subsample, for each LGN and composed the average map and qT1 threshold as above, with the exception that the components were tagged as M and P based on the difference in their mean qT1 values rather than their estimated proportions. The reason for this change was to be able to use the proportion as a measure for the match with the analysis on the average of all 16 maps which showed the mean qT1 values in accordance with the proportions of the M and P regions (higher qT1 and lower proportion vs lower qT1 higher proportion, respectively), but this may not be the case for different subsamples since the mixture Gaussian model may fail to fit to the noisier data. Crucially, we measured the classification accuracy by calculating the percent of match in the categorization of voxels into M and P regions with the subsample vs the entire sample of the 16 maps averaged together.

#### 2.2.3. Functional MRI

fMRI data were acquired over the whole brain with a multi-band EPI sequence with 84 interleaved transversal slices at 1.5 mm isotropic voxel resolution (TR = 1500 ms, TE = 39 ms; α = 75°; FoV= 192 mm, bandwidth = 1562 Hz/Px, phase encoding = A → P), and a slice acceleration factor of 6.

##### 2.2.3.1. LGN Localizer

For each of ten 5-min scans, participants were instructed to fixate on the dot at the screen center. As shown in Figure 3a, a 5 s fixation screen was followed by the 16 s visual stimulus that alternated between left and right hemifields with a 5 s blank between alternations. The initial hemifield was counterbalanced across blocks. Stimuli were presented on a 32-inch LCD BOLDscreen (Cambridge Research Systems Ltd.) with a 60 Hz refresh rate and 1920 × 1080 resolution. The stimuli were prepared and presented in MATLAB using Psychophysics Toolbox (Brainard, 1997; Kleiner et al., 2007; Pelli, 1997) on a Windows computer. The visual stimulus was a black and white checkerboard, hemifield radius of 8.5°, flicking at 4 Hz on a neutral gray background. The fixation point was drawn within a central gap of 0.25° in radius.

**Figure 3.**
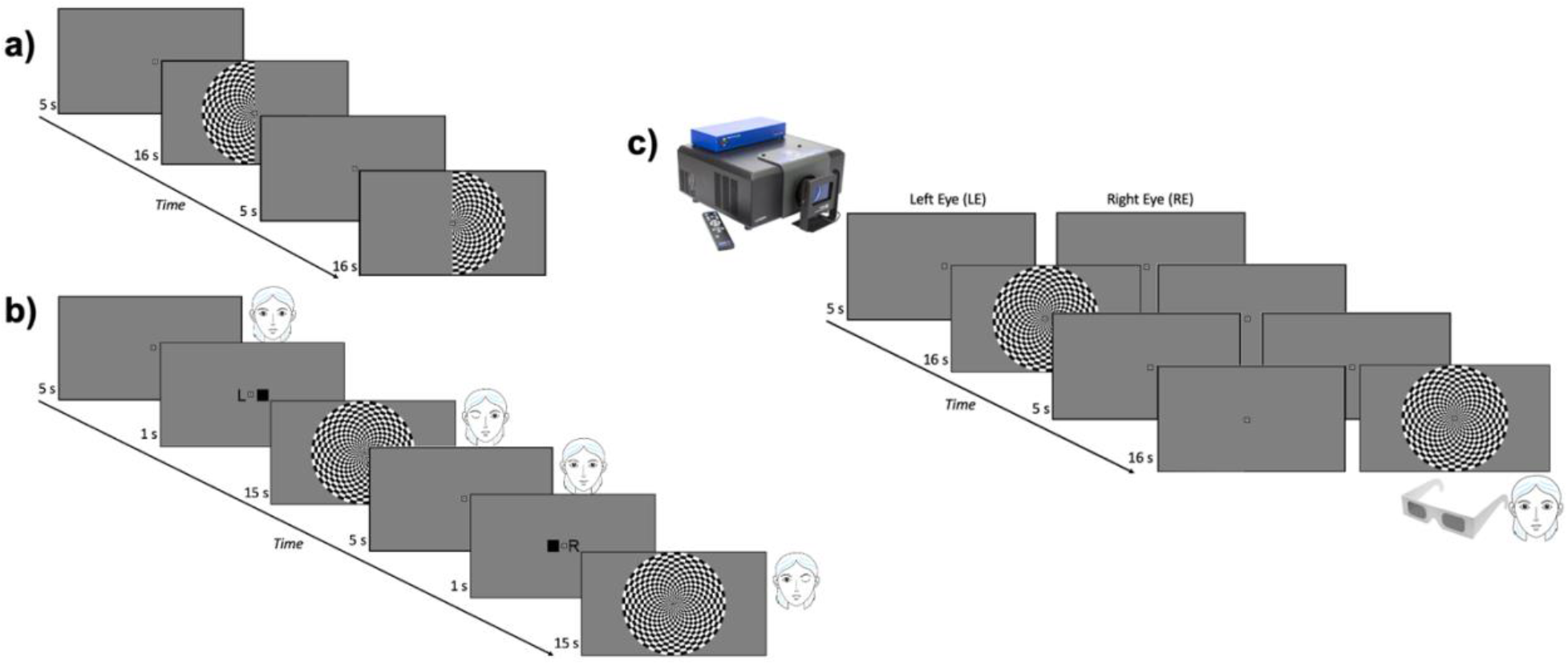
Timeline of fMRI tasks. **a)** LGN localizer with visual hemifield stimulation. **b)** Monocular eye localizer: each eye was stimulated alternately with the other eye closed. **c)** Dichoptic eye localizer: each eye was stimulated alternately with the other eye viewing a neutral gray blank screen.

##### 2.2.3.2. Monocular Eye Localizer

For each of ten 5-min blocks (nine blocks for S2), participants were instructed to fixate on the central dot on the screen, and close one eye at a time when cued. A blank fixation screen was presented for 5 s followed by the instruction: the letter L (respectively, R) on the left (right) side of the central dot and a black square on the right (left) to indicate that the left (right) eye should be open and the right (left) eye closed (Figure 3b). After 1s, the full-field 4 Hz flickering checkerboard (17° diameter, 0.5° central gap) appeared for 15 s while the instruction remained in the central gap. The eye open conditions alternated regularly in a 5-minute block, with the order counterbalanced across different blocks. The software and the materials to prepare and present the stimuli was the same as those for the LGN localizer stimuli (see 2.2.3.1).

##### 2.2.3.3. Dichoptic Eye Localizer

Participants wore circularly polarized paper glasses, and stimuli were presented with a ProPixx (VPixx, Inc.) projector with a 120 Hz refresh rate, 1920 × 1080 resolution, and a circularly polarizing filter in front of the projector lens, synchronized to the frame rate, which allowed for dichoptic viewing at 60 Hz. As Figure 3c illustrates, the timeline of this task was the same as that of the LGN localizer, but with a full-field visual stimulus shown either to the left eye or to the right eye while the other eye was shown the neutral gray fixation screen. The flickering checkerboard stimulus was the same as in the monocular eye localizer task, except 12.5° in diameter, due to the different screen size. The stimuli were prepared using the DataPixx toolbox and Psychophysics toolbox in MATLAB, running on a Linux computer.

##### 2.2.3.4. Data Processing

To pre-process the functional data, we applied intensity normalization, high-pass temporal filtering and motion correction using MCFLIRT. For the binocular LGN localizer only, the data were spatially smoothed with a 2.5 mm FWHM kernel.

The data were analyzed with a generalized linear model (GLM) with two explanatory variables (EVs) for each experiment: left (LH) and right hemifield (RH) for the LGN localizer and left (LE) and right eye (RE) for the two eye localizer experiments. Also, a confound variable was added to the model for the motion outlier volumes, as determined by the fsl_motion_outliers command thresholded at the 75^th^ percentile + 1.5 times the interquartile range. All possible contrasts were computed between the two main EVs. The significance threshold for the LGN localizer was corrected for multiple comparisons using cluster correction whereas no correction was applied for the eye localizer tasks, as they were analyzed in the LGN region of interest defined by the localizer scans. Finally, we conducted a fixed-effects analysis for each participant to combine the multiple scanning runs from each task separately and in combination across tasks.

Before analyzing the eye-specific signals, we first adjusted the LGN masks based on the LGN localizer results. For each subject, we examined the whole brain activity for LH vs RH and RH vs LH to identify the right and left LGN respectively. We outlined the significant activity in the LGN using the LGN mask created from the anatomical qT1 map as an anchor to separate the activity from the adjacent pulvinar (Figure 4c). In some slices, the functional activity was not entirely aligned with the anatomical LGN mask (e.g., S3 in Figure 3c), perhaps due to uncorrected EPI distortions. In this case, the functional activity in that slice took precedence and the boundaries were determined utilizing other cues such as anatomical proportions of the LGN.

**Figure 4.**
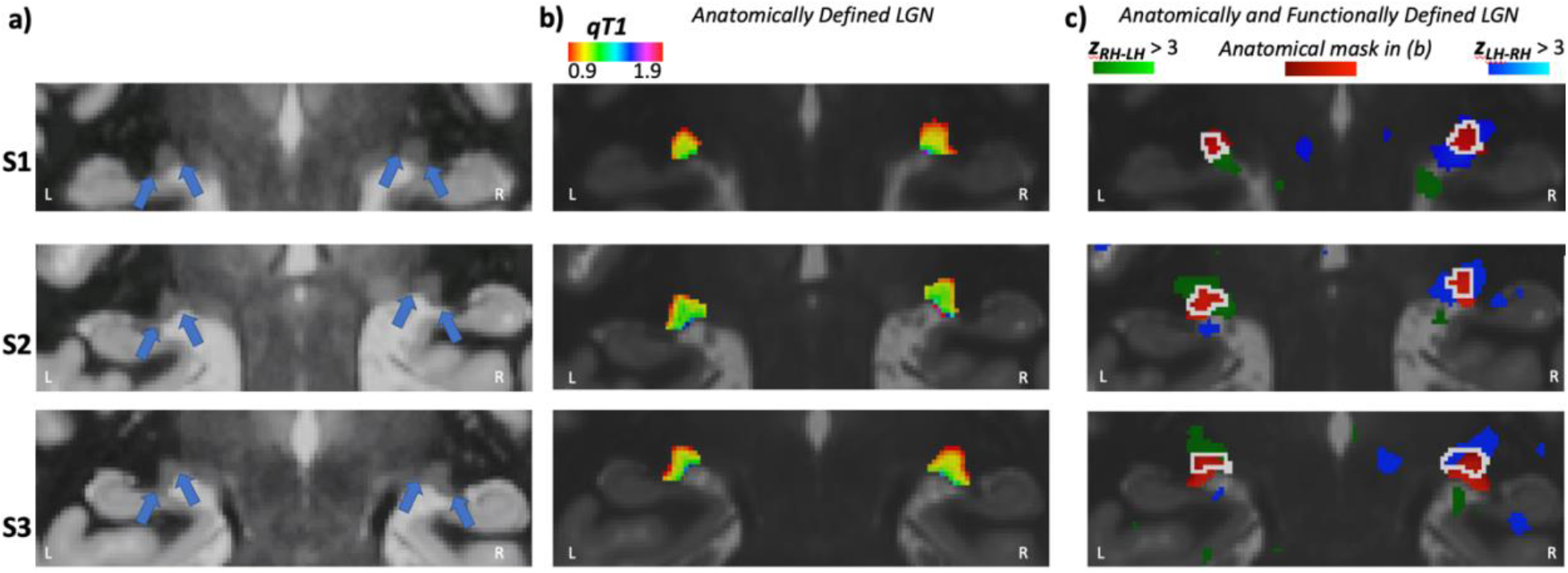
qT1 maps for each participant in separate rows. a) The average qT1 maps of a coronal slice for each subject. Contrast and brightness of the images were adjusted for this illustration. b) The color-coded qT1 maps within each LGN. c) LGN localizer results showing the significant functional activity for the contralateral visual hemifield compared to the ipsilateral hemifield (LH: left hemifield, RH: right hemifield), which is used to adjust the LGN masks for visual activity (white outline).

To identify the eye dominance signals in each LGN, we analyzed the functional data from the monocular and the dichoptic eye localizer tasks. For each voxel in the LGN, we determined its ocular preference based on the *t*-score for the LE vs. RE contrast (Haynes et al., 2005; Qian et al., 2020). A positive *t*-score in a voxel indicated a stronger LE response whereas a negative value indicated a stronger RE response.

## 3. Results

### 3.1. LGN Volume

We calculated the LGN volume for each participant using the LGN masks created from the anatomical qT1 map (Figure 4b). The volume of the left LGN was 138.2, 132.4, and 131.0 mm^3^ for each of three subjects and was smaller than their right LGN, measured as 147.2, 149.6, and 132.4 mm^3^ respectively. We also calculated the LGN volumes using the LGN masks that were adjusted for the significant visual activity (Figure 4c). These functionally adjusted LGN masks resulted in volumes of 125.9, 124.9, and 116.6 mm^3^ for the left LGN and 154.0. 114.9, 130.3 mm^3^ for the right LGN, for each subject respectively. These volumes are smaller than the volumes that were calculated from the anatomically defined LGN, except for S1’s right LGN. All volumes were within the range of 91–157 mm^3^ reported in a histology study (Andrews et al., 1997).

### 3.2. M and P Segmentation with qMRI

The qT1 results for the M and P subdivisions were anatomically reliable. First, as can be seen in the histograms in Figure 5a, the P voxels had shorter T1 relaxation than the M voxels, suggesting more myelination in the P region. The mean qT1 of the identified P voxels were 1.03 ± .003, 1.08 ± .003, 1.03 ± .004 s for left LGN and 1.01 ± .003, 1.08 ± .003, 1.04 ± .004 s for right LGN while the mean qT1 of the M voxels were 1.24 ± .014, 1.33 ± .018, 1.27 ± .016 s for left LGN and 1.19 ± .012, 1.35 ± .023, 1.31 ± .023 s for right LGN, for each subject respectively. The mean difference in the mean qT1 values for M and P was 235.49 ± 14.9 ms. This result is consistent with Müller-Axt et al.’s (2021) who also found shorter qT1 for the P segment and a difference in qT1 between the M and P around 300 ms using a 7T scanner. Also, as can be seen in Figure 4b and 5b, there was a gradual transition in qT1 from P to M region. P voxels that had a qT1 value closer to the threshold (dashed line in Figure 5a) were also spatially closer to the M set. This gradient nature of qT1 map within LGN was consistent across all slices for all subjects. More importantly, the M and P subdivisions conformed to the expected anatomical locations. As evident in Figure 5b, the M voxels occupied the ventromedial region while the P voxels occupied the dorsolateral region of LGN for all subjects. Last, the identified M voxels made up 11.7%, 15.8%, and 16% of the left LGN and 18.9%, 17%, and 16.6% of the right LGN for each subject respectively. These proportions are similar to what had been reported in histology studies (Andrews et al., 1997; Selemon & Begović, 2007). All these results suggest that our M and P segmentation based on the qT1 values in LGN is anatomically consistent in their proportion, myelination, and spatial location within LGN.

**Figure 5.**
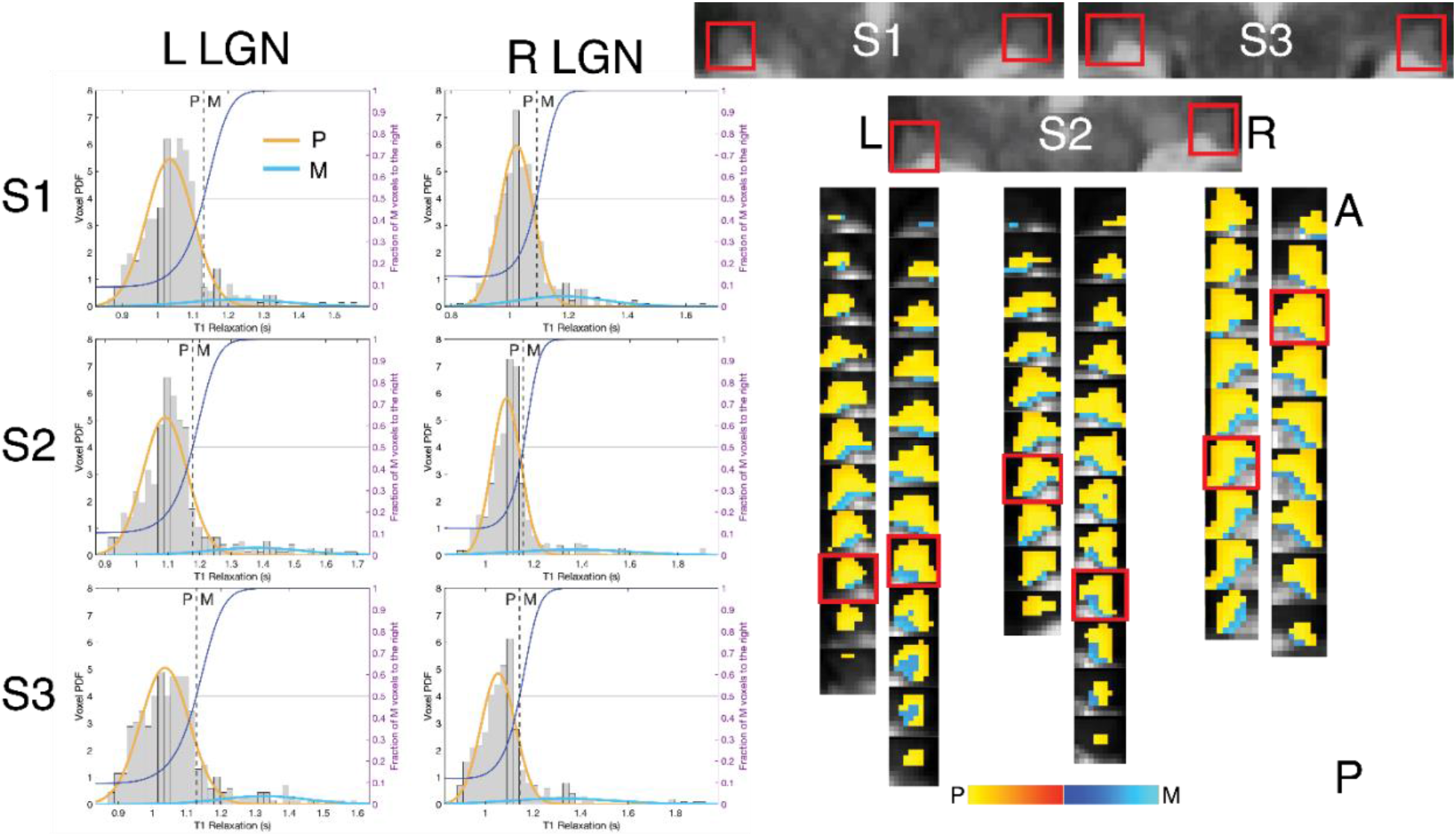
qT1 results for each participant. a) Histograms of voxels for each LGN. The *x*-axis is the qT1 values, the left *y*-axis is the voxel probability density function, and the right *y*-axis is the fraction of M voxels to the right of the distribution for a given cut-off value which is plotted with the dark blue line. Yellow and blue lines show the Gaussian fits for M and P resulting from a two-component mixture model of the data while the dashed line shows the actual cut-off for M and P where the fraction of voxels belonging to the right of the distribution is 50%. b) qT1 maps color-coded within the M and P, separated based on the cut-off indicated by the dashed line in a. The red outlined are the slices corresponding to the reference coronal slice for each LGN.

We acquired more qT1 data (16 volumes per subject) than were necessary, to determine how much data was required at 3T to obtain reliable results. We randomly subsampled the set of qT1 maps and repeated the M and P segmentation process for different numbers of volumes used for each average map as well as for each volume separately. In general, only a small number of averages were required to reliably produce the same result as averaging 16 volumes. Figure 6 illustrates the match between the subsample and the entire sample, in terms of the percent of voxels classified as the same (M or P) with the qT1 analysis. The blue dots represent each subsample and there were 16 dots for each subsample size on the *x*-axis, except the entire sample of 16 maps. The Gaussian mixture model generally failed to fit the data for the individual maps (*N* = 1), resulting in a very low classification accuracy. The black bars show the mean match for each subsample size on the *x*-axis. Fitting an exponential shows convergence to an asymptote (dashed line) at 95.53% consistent categorization on average. The number of volumes to reach 95% of this asymptote (lower edge of the shaded area around the dashed line) ranged from 2–5, corresponding to 1–1.5 hours of scanning. We also calculated the proportion of M in the entire LGN as a function of the number of averaged maps in each subsample (see Figure S1 in the supplemental). This measure also agreed with the conclusion that 1–1.5 hours of data were sufficient for reliable segregation of the M and P regions in the LGN.

**Figure 6.**
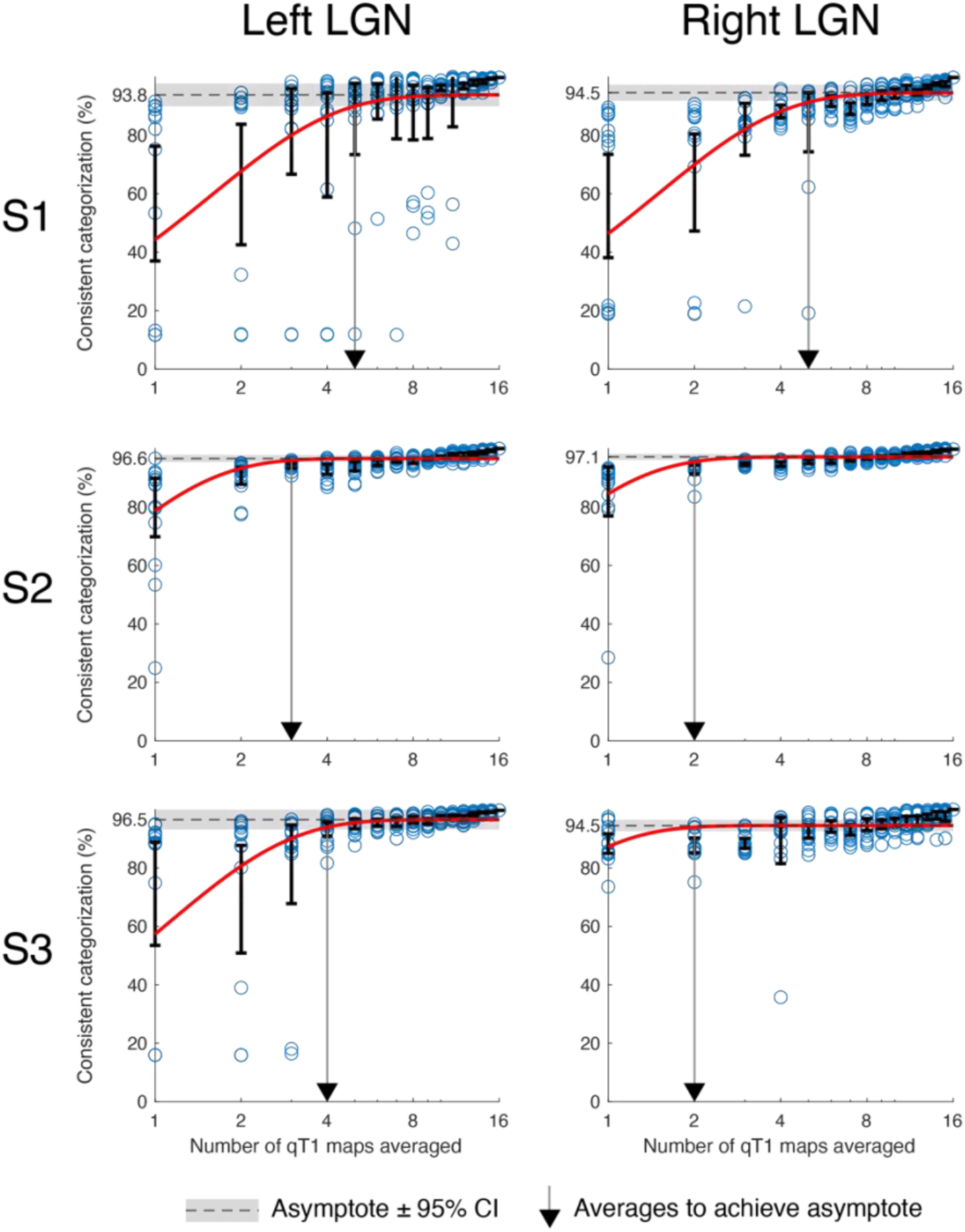
Results for the random subsampling of qT1 maps for each LGN. On the *y*-axis is the percent of voxels classified as the same with the subsample vs the entire sample. On the *x*-axis is the number of qT1 maps used in the subsample. Each blue dot represents a subsample while the dot on *x*=16 represents the entire sample. The black bars are centered around the means of 16 subsamples of different sizes (i.e., 16 blue dots for *x*=1 through *x*=15); error bars indicating 95% confidence intervals (CIs). The red curve is the exponential growth curve fitted on the log of the means, using the formula 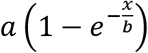, where x is the subsample size on the x-axis; a and b are the estimated parameters where a is the percent at which the curve reaches to a horizontal asymptote which is indicated by a dashed line with the shades indicating the 95% CIs. The vertical gray line is where the fitted curve reaches the lower bound of the asymptote.

### 3.3. Eye-specific Segmentation with fMRI

The results of the monocular and dichoptic tasks, as well as a statistical combination of the two tasks, are shown in Figure 7 for a representative subject. However, the results were similar for the other subjects (see Figure S1 in the Supplemental Materials). Figure 7 color codes the voxels as responding to the contralateral (blue) and ipsilateral (red) stimuli for each LGN when calculated based on the sign of the *t*-score for LE vs RE contrast. The monocular task resulted in a stronger ocular preference compared to the dichoptic task. This result can be seen in Figure 7 in the combined results for the two tasks (third row) which appeared more similar to the monocular condition. Accordingly, the dichoptic task significantly activated fewer voxels (Figure 7 bottom). Thus, the eye signals were stronger in LGN when the other eye was closed instead of being open and presented with a blank screen. The contribution from the non-stimulated eye on the signals for the stimulated eye differed between tasks.

**Figure 7.**
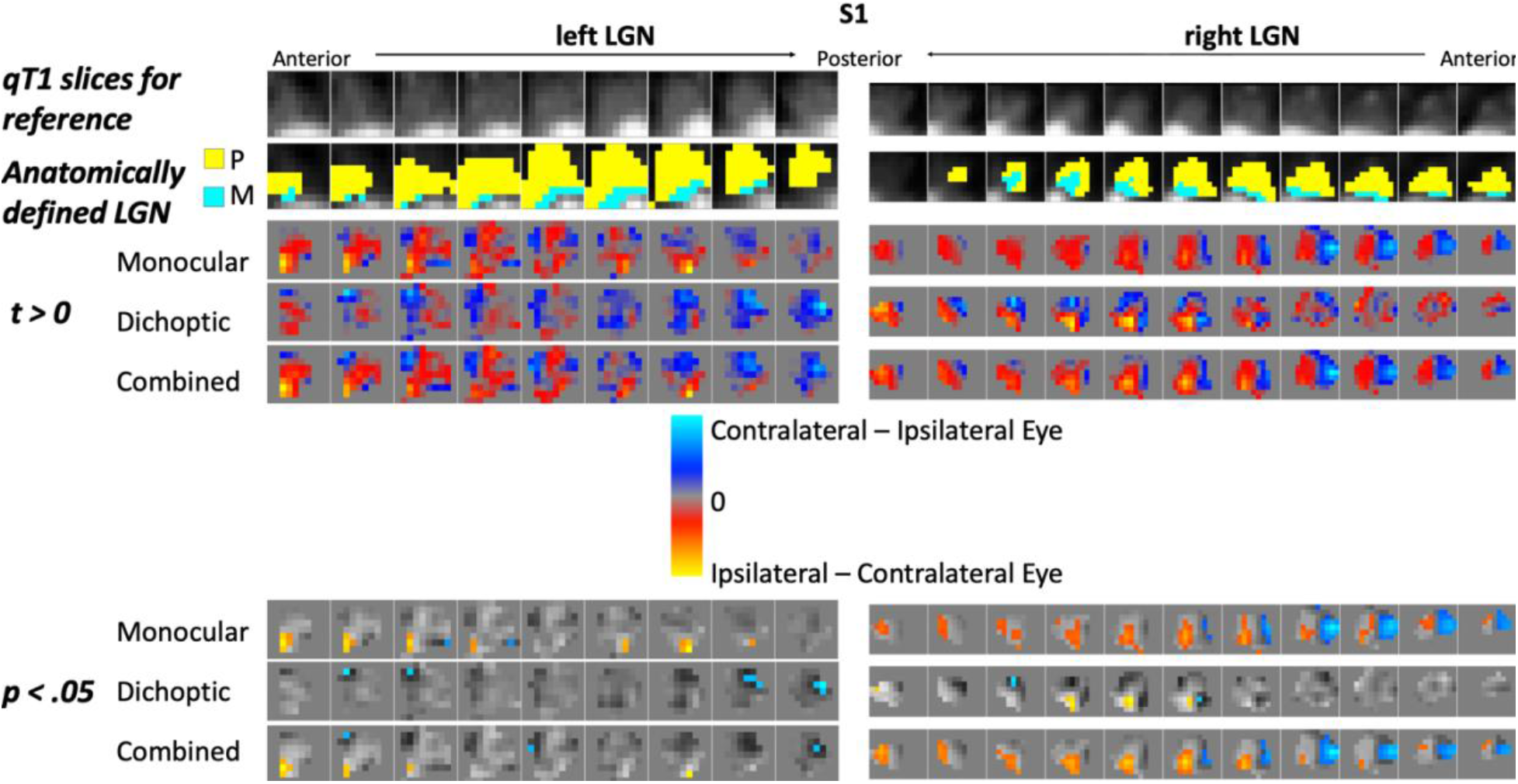
Eye-specific activity for a representative participant (S1). For reference, the average qT1 map and the anatomically defined LGN (Figure 4b) with the M and P segregation based on the qT1 analysis was shown on the first two rows for the same slices as in the images below them. The LGN masks for the eye-specific fMRI results were anatomically and functionally identified as illustrated in Figure 4c. For the monocular eye localizer, participants closed one eye at a time. For the dichoptic eye localizer, one eye was shown blank while the other eye was visually stimulated. The ocular preference was calculated based on the *t* values for Left Eye > Right Eye in the eye localizer analysis, changed here to show the contralateral and ipsilateral eye for illustration. On the bottom are the voxels showing significant ocular preference for Left Eye > Right Eye contrast.

There was a RE dominance evident in both tasks. As seen in Figure 7, there were more ipsilateral voxels in the right LGN (red) while more contralateral voxels were identified for the left LGN (blue). The exceptions to this RE bias were observed in the monocular task in S1’s left LGN, which had more voxels preferring the LE, and in S3’s left LGN, which had equal number of RE and LE voxels (also see Table 1). On average, the percentage of RE voxels to the LGN was 56.5% for the left LGN and 65% for the right LGN with the dichoptic task whereas it was 44.5% for the left LGN and 64.3% for the right LGN with the monocular task.

**Table 1.**
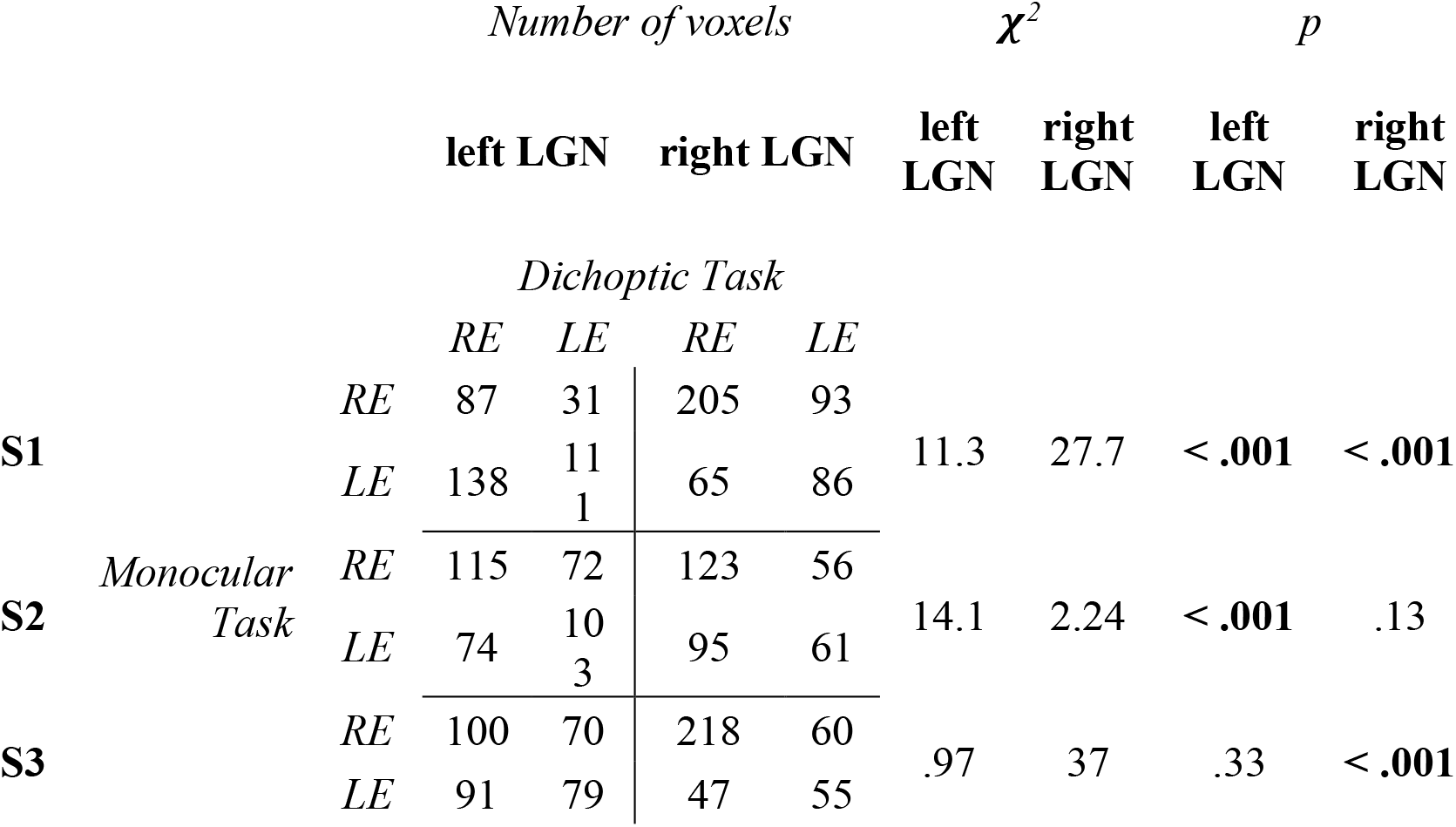
Chi-square results for Left Eye (LE) and Right Eye (RE) categorization in each LGN

Figure 8 displays the correlations for each voxel between the ocular preferences with different tasks, i.e., *t*-score for LE vs RE. The correlation between the tasks was significant for five out of six LGN, *p’s* ≤ .001. The LGN that did not show significant correlation (S2 right LGN) also failed to show voxels that were significant in their ocular preference in the combined analysis of the two tasks, as indicated by the red dots in Figure 8. All the significant correlations increased when we used only the significant voxels from the combined analysis (the third row in Figure 7 and Figure S1).

**Figure 8.**
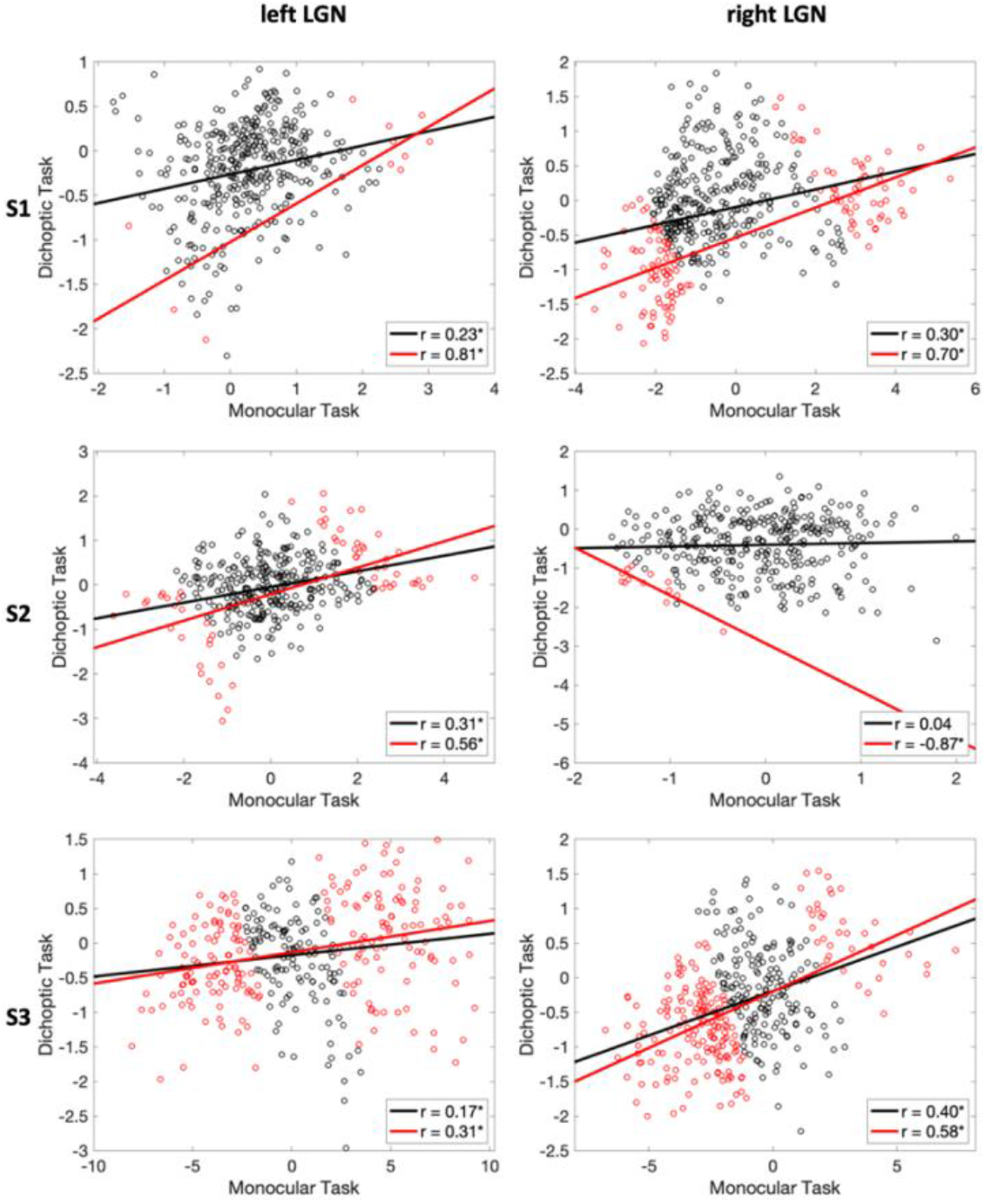
Scatterplots of voxels showing the ocular preference with the monocular and dichoptic tasks for each participant. The red dots are the voxels whose ocular preference was significant in the combined analysis of the two tasks. The solid lines show the correlation between the two tasks (black for all voxels and red for only the significant voxels. \**p* ≤ .002

To test the significance of the LE vs. RE classification based only on the sign of their *t*-scores and ignoring the magnitude, we conducted a chi-square analysis for each LGN on the resulting categorical variables (Table 1). The classification of the voxels matched between the two eye localizer tasks on only four of the six LGN, *p*’s < .001. These significant results were driven by the RE bias in the classification. A close inspection of the cross-tables in Table 1 indicated that there was more match between the RE voxels (the upper left cell for each subject’s each LGN) than between the LE voxels (the lower right cell) for the majority of the LGN. Table 2 shows the chi-square results for the voxels showing significant ocular preference in the combined analysis of the two tasks. The right eye bias was reduced when only the significant voxels were classified and the classification with the two tasks significantly matched again for four out of six LGN, *p*’s < .001.

**Table 2.**
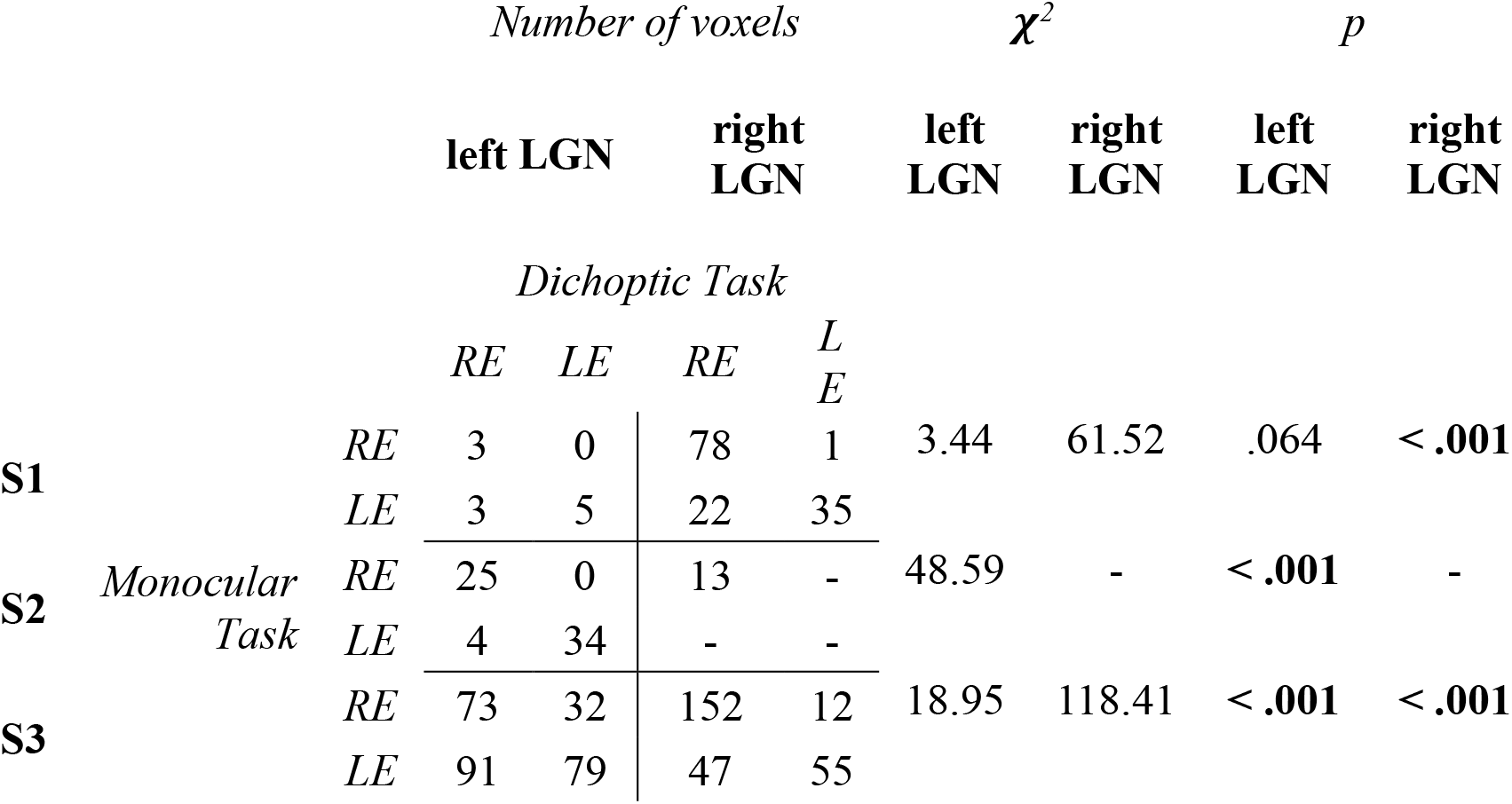
Chi-square results for Left Eye (LE) and Right Eye (RE) categorization in each LGN for only the voxels that showed significant ocular preference in the combined analysis.

Our goal in identifying the eye-specific regions was to segment the M and P layers. First, we could not quantitatively compare the results from the qMRI processing with the fMRI processing because of the mismatch between the LGN masks when adjusted for functional activation (see Section 2.2.3.4). More importantly, we tried to identify the contralateral layers positioned most ventrally or dorsally to find the contralateral M or P layers, respectively. The dorsal contralateral region appeared robustly with either eye localizer task for all LGN using the signed classification, and in four of six LGN using only the voxels activated significantly by the combination of both tasks (see Figure S1). However, the ventral contralateral layer did not appear reliably. We could identify the ventral contralateral layer in only two LGN in one or other of the tasks, though the successful task differed between the two LGN (monocular task for S1 left LGN and dichoptic task for S3 left LGN; see Figure S1). Using only the significant voxels in the combined analysis (Figure S1, third row for each subject), the ventral contralateral layer could only be identified for S3. Instead of reliable individual layers, we could identify a contralateral eye cluster located more dorsolateral and an ipsilateral eye cluster located more medioventral.

## 4. Discussion

To segregate the M and P regions in human LGN, we used MRI methods that were not dependent on the stimulus characteristics, unlike the previous attempts (Denison et al., 2014; P. Zhang et al., 2015) that were confounded by the stronger activation of hilum of LGN (DeSimone & Schneider, 2019). Using qT1 (i.e., measuring the T1 relaxation time for each voxel), we successfully identified the M and P components of both LGN in all the subjects, which conformed to our anatomical expectations. However, attempting to identify the individual ipsi- or contralateral layers using fMRI was less successful. The identification of the eye-specific regions was more consistent with the monocular task than the dichoptic task. The P layers in the dorsal contralateral cluster could be readily identified, but the ventral contralateral (M) layer was not consistently activated.

qMRI, with a MP2RAGE sequence we used, has been shown to be more advantageous for precisely imaging subcortical structures. Aldusary et al. (2019) compared different T1 sequences for LGN volume and found that MPRAGE imaging was more accurate compared to proton density imaging with a 3T scanner. Using the MP2RAGE sequence with two inversion times allowed us to calculate the T1 parameter for each voxel, which enhanced the segmentation of the whole LGN relative to an MPRAGE or proton-density weighted sequence and allowed the segmentation of the M and P divisions. The whole LGN volumes we found were consistent with post-mortem histology (Andrews et al., 1997) and with the structural MRI studies with a 3T scanner (proton density imaging in Giraldo-Chica et al., 2015; phase difference enhanced imaging in Kitajima et al., 2015; T1-weighted imaging in Wang et al., 2015). On the other hand, when defined functionally, previous studies reported much higher volumes of LGN (Denison et al., 2014; Kastner et al., 2004), likely as a result from the difficulty of segmenting the LGN from surrounding visually active regions such as the lateral and medial pulvinar.

To segregate the M and P regions based on the qT1 maps, we followed a data-driven approach to replicate Müller-Axt et al. (2021) at 3T. By fitting a two-component model to the qT1 data, we selected the smaller proportion component as M and the larger proportion component as P. The P component showed shorter T1 relaxation time (i.e., qT1) than the M component, indicating more myelination in the P region. This is consistent with Müller-Axt et al.’s (2021) results (also see preprint Oishi et al., 2020) and with higher cell density and more myelination in the P compared to the M divisions (Hassler, 1966; Pistorio et al., 2006). Previous studies used a fixed proportion as the criterion to segregate the M and P sections (Denison et al., 2014; Oishi et al., 2020), based on the histology findings that, on average, 20% of the LGN is M (Andrews et al., 1997; Selemon & Begović, 2007), but this approach, even if correct, would not allow the independent measurement of the M division properties. Individuals show great variation in the proportions of the subdivisions (Andrews et al., 1997; Müller-Axt et al., 2021), and indeed our participants had M divisions ranging from 12–19% of the LGN volume. We were able to identify the M and P subdivisions in individual subjects, compared to in a group as was shown in Müller-Axt et al. (2021). We acquired a large amount of data in a small number of subjects to perform a subsampling analysis to demonstrate that 1–1.5 hours of scanning at 3T was sufficient to reliably classify the M and P voxels in individual LGN.

In the thalamus and brainstem, pulsatile motions are a concern that can cause noise in the images. We were nonetheless able to obtain reliable results with the qT1 scans after averaging as few as 2–5 volumes. Also, obtaining reliable qT1 measurements in the M region requires that the bright cerebrospinal fluid (CSF) be excluded from the LGN masks (McNab et al., 2013). Upsampling the qT1 image helped reduce partial volume effects and exclude the CSF from the LGN masks. Also, the thickness of the M region (Figure 5b) relative to the voxel size (0.7 mm isotropic) was sufficient to reliably distinguish the CSF from M, enabling us to exclude the CSF as a significant contaminating factor to the qT1 estimates for the M region. Similarly, the hilum could potentially be mistaken for the M section, as blood vessels show T1 values similar to M region (X. Zhang et al., 2013). However, Figure 5b indicates that the geometry of the M region was not consistent with a hilum confound, i.e., the identified M region did not generally intrude dorsolaterally into the interior of the LGN as would the hilum, except perhaps in two posterior slices in one subject. Given that this intrusion was rare, and combined with the lack of any observed hilar structure in Müller-Axt et al.’s (2021), we conclude that the interior intrusion of the M subdivision in these two slices was mostly likely a result of the folding structure of the LGN and not the hilum.

Our investigation of the eye dominance signals did not yield consistent results with the monocular vs. dichoptic eye localizer tasks when analyzed with a generalized linear model (GLM). We found that the eye-specific signal amplitudes were larger with the monocular than dichoptic task such that it was difficult even to measure significant activation with the dichoptic task despite the same amount of data. This might indicate interference or rivalry from the “non-stimulated” eye during dichoptic presentation. Both tasks revealed right eye dominance in all three subjects, comprising 57.6% of the LGN volume on average. The classification of the RE voxels was more consistent between the tasks than for the LE voxels. Previous studies identifying voxel eye preference used these two tasks separately (Haynes et al., 2005; Qian et al., 2020). Using the dichoptic task, Qian et al. (2020) identified significant eye-specific clusters with GLM at 7T. Here, we show that the GLM was not suitable to detect significant eye-specific activations at 3T with dichoptic presentation. This poses a problem because dichoptic presentation can be coded by the experimenter to control which eye to be stimulated while the monocular task requires subjects to close each eye alternately that cannot be controlled by the experimenter unless an eye tracking device is used.

In our investigation of the M and P eye signals, we could not compare the results from fMRI with qMRI as there was not an exact match between the voxels of the LGN masks used for the two. Adjustment of the anatomical LGN masks for the visual activity was not expected to make such a difference; however, this is well beyond our study and perhaps related to the EPI distortions. Crucially for the eye-specific signal investigation though, we found that the dorsal contralateral-eye region, classified as P, could be reliably identified with both monocular and dichoptic tasks, whereas the ventral contralateral-eye layer, which would be classified as M, could not be reliably activated with either task. The fMRI resolution we used (1.5 mm isotropic) is not optimal for imaging the contralateral M layer, and it is difficult to improve this resolution without the signal being lost in the noise at 3T. Other techniques for thin layer segmentation could be used such as anisotropic voxels with slices parallel to the LGN (e.g., Kashyap et al., 2018), although this requires subjects with a particular LGN geometry and perhaps unconventional positioning in the scanner and enhanced motion suppression. Critically for our fMRI methods, the hilum region of LGN did not dominate the responses to eye-specific stimuli when the other eye was closed. However, the right eye bias did interfere with our ability to classify the eye-specific layers, and functionally identifying the eye-specific layers does not appear to be a promising approach to segmenting the M and P regions of the LGN.

In summary, our qT1 results using a 3T MRI scanner replicated measurements performed at 7T (Müller-Axt et al., 2021). Our results at the individual subjects indicate that this qMRI method and analysis can be used for M and P segmentation with only 1.5 hours of data, a reasonable application time for clinical and research purposes. Our fMRI results for eye-specific region segmentation using GLM were much more reliable when subjects closed an eye (monocular stimulation) than when stimulating only one eye with both eyes open (dichoptic).

## Acknowledgments

This work was supported by the National Institutes of Health (NIH/NEI 1R01EY028266 to KAS).

We would like to thank Ibrahim Malik, Joy Lin, Christina Nelson, Jack Melchiorre, Anton Lebed, and Heather Aiken for their assistance during data collection.

## Notes

### Competing Interest Statement

The authors have declared no competing interest.

