## Supplemental Figures for "Evaluating Quantitative and Functional MRI As Potential Techniques to Identify the Subdivisions in the Human Lateral Geniculate Nucleus"

### Supplemental Materials

| | *AIC* | | | | *BIC* | | | |
|---|---|---|---|---|---|---|---|---|
| | **left LGN** | | **right LGN** | | **left LGN** | | **right LGN** | |
| | *Number of components in the GM* | | | | *Number of components in the GM* | | | |
| | *1* | *2* | *1* | *2* | *1* | *2* | *1* | *2* |
| **S1** | -758.58 | -849.02 | -809.5 | -950.95 | -750.58 | -829.02 | -801.38 | -930.64 |
| **S2** | -534.88 | -701.9 | -478.41 | -843.98 | -526.97 | -682.12 | -470.25 | -823.59 |
| **S3** | -561.31 | -698.17 | -432.72 | -650.16 | -553.41 | -678.44 | -424.8 | -630.38 |

**Table S1.** Akaiken Information Criteria (AIC) and Bayesian Information Criteria (BIC) for the 1-component and 2-component Gaussian Model (GM) on the average qT1 map of each LGN.

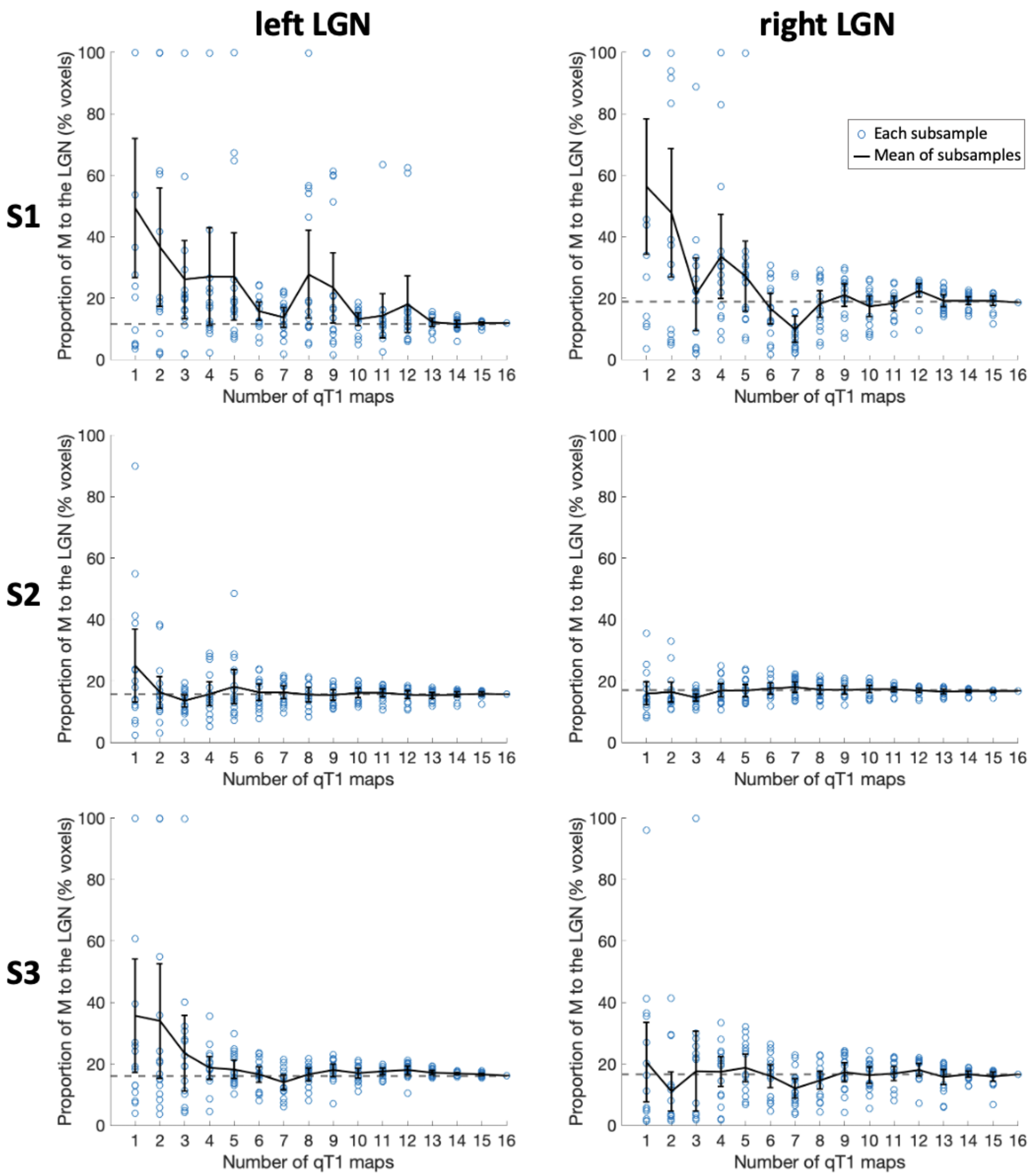


**Figure S1.** Results for the random subsampling of qT1 maps for each LGN. On the *y*-axis is the proportion of M to the LGN at the threshold qT1 value found by the qT1 analysis. On the *x*-axis is the number of qT1 maps used in the subsample. Each dot represents a subsample while the dot on *x*=16 represents the entire sample whose proportion of M is indicated with a dashed line (calculated based on the dashed line in Figure 5a). The black line is the mean of 16 subsamples of different sizes (i.e., 16 blue dots for *x*=1 through *x*=15); error bars are 95% confidence intervals.

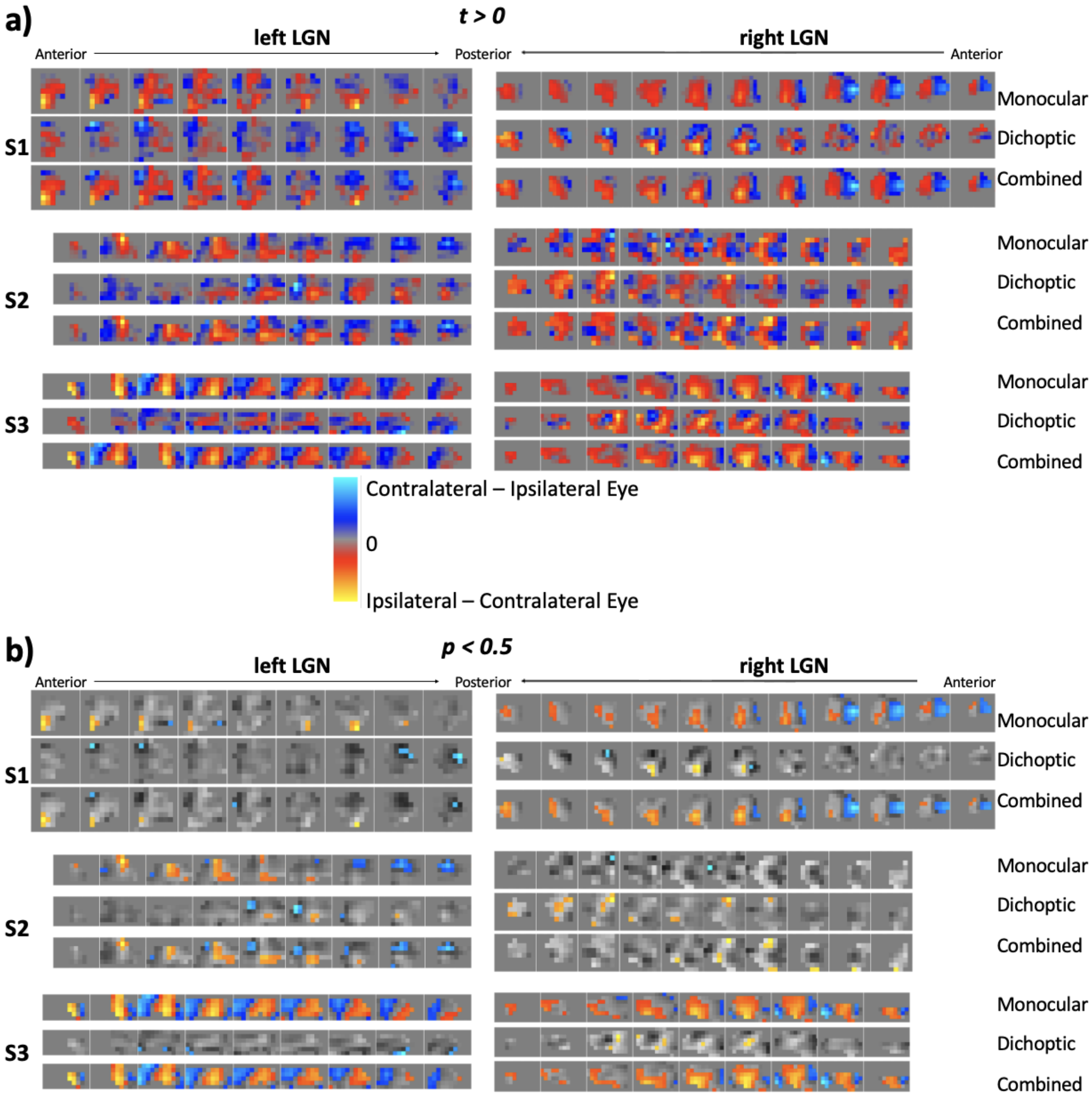


**Figure S2.** Eye-specific activity for each participant's lateral geniculate nucleus. For the monocular eye localizer, participants closed one eye at a time alternately. For the dichoptic eye localizer, one eye was shown with blank while the other eye was visually stimulated alternately. a) The ocular preference was calculated based on the *t* values for Left Eye > Right Eye in the eye localizer analysis. Here, these *t* values were converted to show the contralateral vs ipsilateral eye preference. b) The voxels showing significant ocular preference in a, uncorrected for multiple comparisons for the voxels.
